# Strain and sex effects on blood accumulation of lead, chromium, and cadmium in strains from the Collaborative Cross mouse population

**DOI:** 10.1101/2025.10.27.684896

**Authors:** Brittni Ming-Whitfield, Danila Cuomo, David W Threadgill

## Abstract

Lead (Pb), chromium (Cr), and cadmium (Cd) are heavy metals that contaminate sites throughout North America. Historically, toxicological effects of Pb, Cr, or Cd compounds have been investigated in a hybrid mouse strain, B6C3F1. However, humans have more genetic diversity and population variability in response to toxicants than is represented in this homogeneous mouse model, which leaves genetic effects on dose response uncertain. Use of the Collaborative Cross (CC) addresses the problem of limited genetic diversity inherent in models like B6C3F1. In previous work, blood Pb levels in panel of female CC lines exposed to high-dose (0.1%) lead acetate showed a strain dependent response. Four strains from the original study with varying Pb blood levels after exposure were selected to determine if strain and sex dependence was exhibited in a two-week acute exposure to Pb, but also to Cr or Cd exposure. To investigate genetic background influence on metal deposition, five animals of each sex from each strain were placed on an American diet for one week prior to dosing high- (0.1%) or low- (0.01%) dose Pb acetate, high- (0.1%) or low- (0.01%) sodium dichromate, or high- (0.1%) or low- (0.01%) cadmium chloride via drinking water ad libitum for 14-days, matching the standard short-term exposure of the National Toxicology Program. Body composition was measured before the start of dosing and prior to necropsy using EcoMRI. Blood Pb at necropsy from this study suggests the strain dependent trends observed in previous exposures is conserved for acute Pb exposure but with different trends for Cr and Cd indicating that even a small panel of strains will not suffice for estimating variation across all toxicants.

## Introduction

Heavy metal pollution in air, water, and food, such as lead (Pb), hexavalent chromium VI (Cr VI), and cadmium (Cd), is an ecological and public health concern. These heavy metals have a variety of adverse health effects and have no safe environmental level. Adverse outcomes are affected by dose, but also by other endogenous and exogenous factors such as genetics and diet (Cuomo et al. 2022). However, sex and dose effects in the context of genetic variation has not been thoroughly explored.

Lead is a toxic, naturally occurring heavy metal in the earth’s crust that can be introduced to the environment through industrial activities. While human action levels of Pb in blood are 3.5 micrograms per deciliter (EPA), there is no safe level of lead at any developmental stage. Children are particularly susceptible to Pb toxicity due to their frequent hand to mouth behavior and developmental stage. Genetic susceptibility to Pb toxicity has been explored in the mouse Collaborative Cross (CC) population (Cuomo et al. 2022) but sex differences remain to be elucidated.

Hexavalent chromium is a toxin present in multiple environmental and occupational contexts. People are typically exposed through particulate matter in air, contaminated ground water, or soil (Thompson et al. 2013). In addition, Cr IV is carcinogenic and greatly increases risk in occupational settings including leather tanning and electroplating. The maximum allowed contaminant level in drinking water is 0.1 mg/L or 100 ppb for total Cr, which is assumed to be Cr IV by regulatory agencies like the EPA. Hexavalent chromium is dangerous due to its creation of reactive oxygen Cr IV toxicity is specific to its oxidative state. The effects of reactive oxygen species from Cr IV can be inhibited significantly by melatonin (Lv et al. 2018); melatonin inhibits Cr induced loss of mitochondria membrane potential. Hexavalent chromium first enters cells in part due to its structural similarity to sulfates and phosphates (Katz and Salem 1993).

Exposure to Cr can cause a variety of adverse effects. Chronic exposure is carcinogenic, while chronic and acute exposures lead to a variety of issues throughout the body including respiratory distress, stomach ulceration, small intestine irritation, and allergic sensitization (Agency for Toxic Substances and Disease Registry (ATSDR) 2012b). Reproductive issues are not thoroughly understood in humans, but abnormal sperm, pregnancy, and childbirth complications have been reported (Thompson et al. 2020).

Cadmium is a heavy metal naturally occurring in the Earth’s crust and is toxic in small amounts. Cadmium is present ubiquitously in rocks, coal, and some mineral fertilizers (Agency for Toxic Substances and Disease Registry (ATSDR) 2012a) as well as in some consumer products such as batteries. It is found in a 2+ oxidative state in nature and unlike Cr, does not undergo reduction or oxidization reactions. Cadmium exposure is most likely to occur in occupations like manufacturing and construction such as metal smelting, refinery, and manufacture of batteries, plastics, solar panels, and coatings (Bernhoft 2013). Most exposure in the US occurs from cigarette smoke, contaminated food consumption, or drinking water. The EPA sets limits for Cd in water at 0.04 mg/L or 40 ppb. Blood levels are indicative of recent exposure while urinary levels can be used for both recent and past Cd exposure.

Research has shown that there is broad susceptibility to a wide range of toxins influenced by genetics (Rusyn et al. 2022). Previous research in our group has elucidated how genetic variability in female CC mice can contribute to variation in blood Pb level accumulation (Cuomo et al. 2022). However, variation in blood levels from exposure has not been investigated for Cr IV and Cd nor in males. This study uses the CC as a model to better elucidate toxicant risk for short-term exposure periods for Pb and the additional heavy metals, Cr IV and Cd.

## Materials and methods

### Experimental design

To understand the strain and sex-specific levels of Pb, Cd, and Cr accumulation in the blood, five male and five female mice from five mouse lines were exposed to 100 ppm or 1000 ppm of lead acetate, sodium dichromate dihydrate, or cadmium disulfide. Blood metal levels were quantified after 14 days of exposure (Fig. 1).

**Fig. 1.**
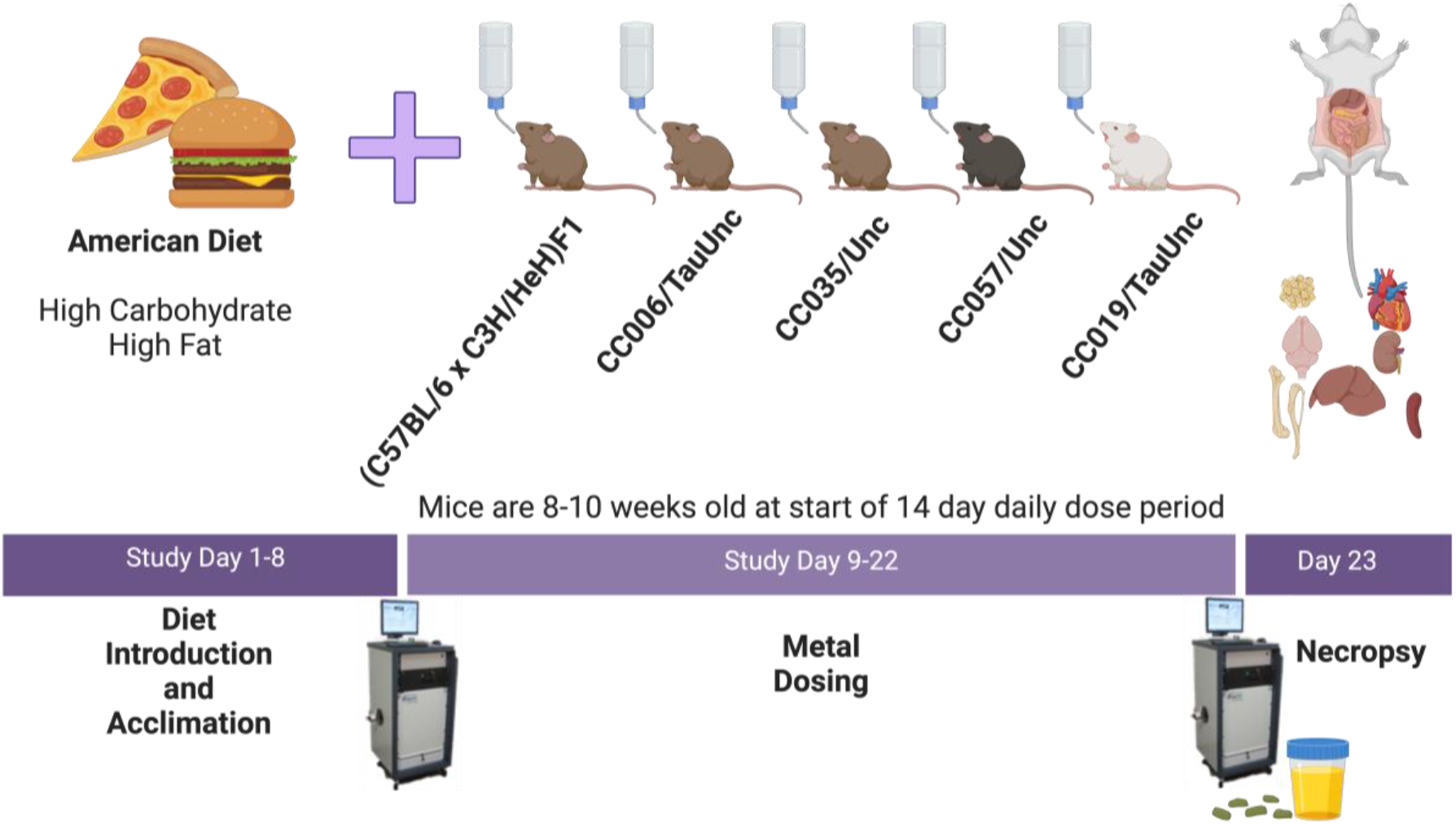
Experimental design to elucidate strain effects on metal levels in blood. At 8-10 weeks of age, mice of each strain and sex were assigned to a low or high dose of each of the three metals. MRI measurements were conducted one day prior to dosing and the day prior to necropsy.

Four CC lines (Churchill et al. 2004; Threadgill et al. 2002) were obtained from an onsite breeding colony at Texas A&M. Strains utilized included CC006/TauUnc (CC006), CC035/TauUnc (CC035), CC057/TauUnc (CC057), CC019/TauUnc (CC019), and B6C3F1/J (B6C3F1) as a control for mice traditionally used in the National Toxicology Program (NTP). Mice were housed in ventilated cages in groups of 2-5 animals. Singly housed animals were avoided unless necessary due to behavioral issues. Mice were fed Standard Diet (Teklad Global Diets, No. 2919) from weaning until the start of the study. Mice were then fed an American diet (protein content 15.8%, fat 16.6%, carbohydrate 53.8%, fiber 2.6%) (Barrington et al. 2018) and were allowed to acclimate to the American diet for 1 week prior to the start of metal dosing. Mice were on American Diet for the entirety of the dosing period and allowed to drink from dosed water sources ad libitum. At the end of the study, mice were euthanized via carbon dioxide asphyxiation and whole blood was collected for metal quantification. Animal use and experiments were approved by the Texas A&M Institution of Animal Care and Use Committee (Protocol 0435-2019).

### Phenotypes

Water consumption was measured throughout the dosing period by water bottle mass and was also measured for a 12 hour-period in metabolic chambers for all mice in the study. Whole blood samples were collected at necropsy via cardiac puncture in 0.5 EDTA tubes and submitted to the Michigan State Veterinary Medical Diagnostic Laboratory (MSUVDL) for quantification of Pb, Cr, and Cd via Inductively Coupled Plasma Mass Spectrometry (ICP-MS) to confirm metal concentrations. All concentrations herein are reported in micrograms/deciliter (ug/dl). Additional phenotypes that may contribute to variance in blood metal levels were measured as well, including body weight before (day 8) and after (day 22) exposure to metals and fat and lean mass before and after exposure using EchoMRI-700. Tissues collected at necropsy included heart, gonadal fat pad, brain, liver, kidneys, spleen, and muscle for flash freezing in liquid nitrogen.

### Statistical Analysis

Statistical analysis was performed using JMP version Pro 17, R statistical software version 4.3.2, and GraphPad Prism. Groups were compared using one way ANOVA with p≤0.5 being significant. Strain, sex, and strain-by-sex effects on metal levels were evaluated using a linear mixed effects model to quantify their impact on blood metal levels. R-output for partitioning of variance is in Supplemental Fig. 1 and Fig. 2.

## Results

### Collaborative Cross Pb accumulation repeatability

CC strains exposed to Pb for 14-days had similar inter-strain variability as strains exposed for 28-days (Fig. 2), although means for each strain were lower than mean blood Pb accumulation recorded after 28-days of exposure (Cuomo et al. 2023). Nonetheless, rank order of blood Pb accumulation was conserved between the studies, with CC006 being the highest and CC019 the lowest Pb accumulator in blood. One-way ANOVA comparisons with Tukey correction showed that CC019 was the only strain without significantly different mean blood Pb accumulation between exposure durations (Fig. 2).

**Fig. 2.**
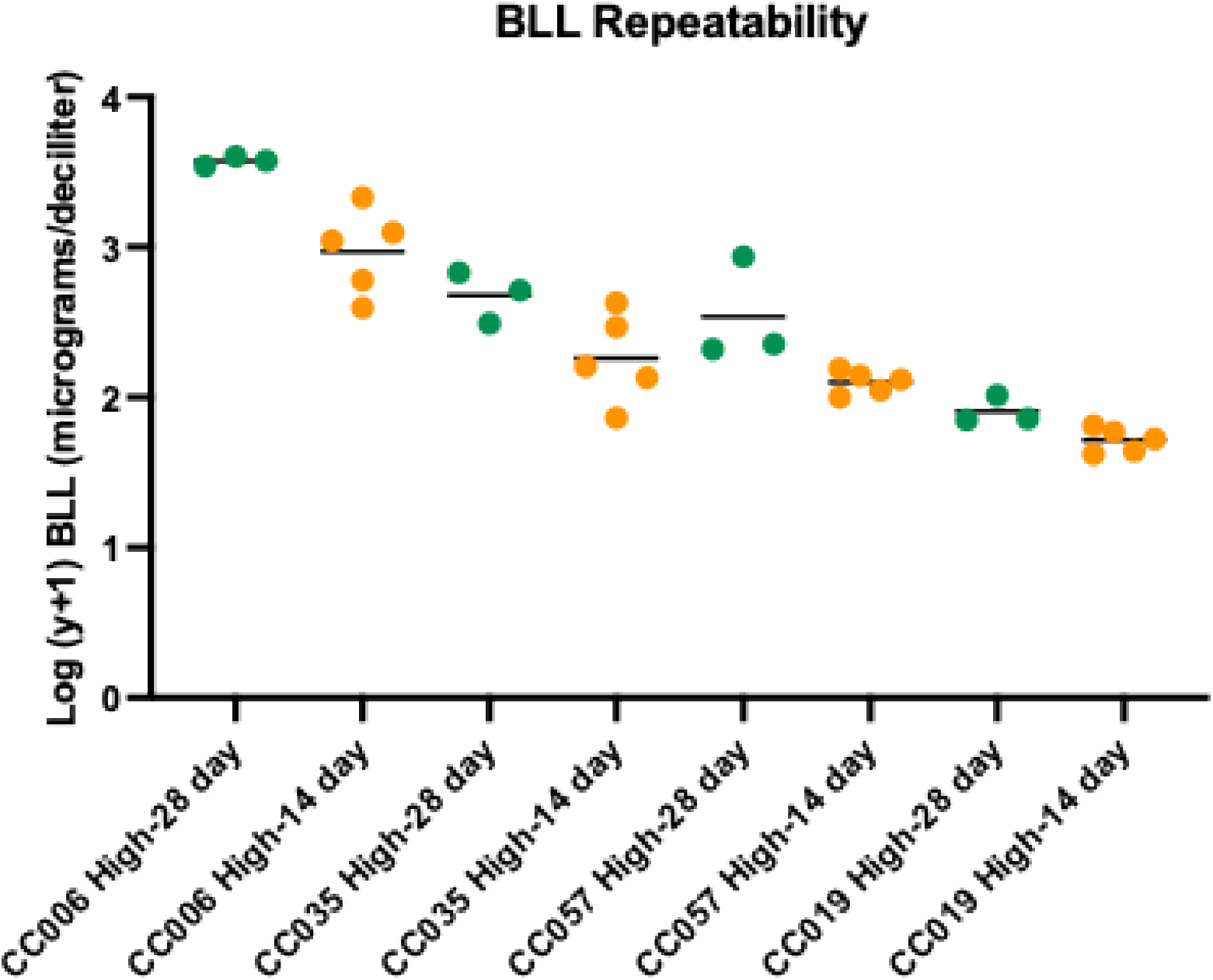
Repeatability of Strain Dependent Blood Lead Accumulation. Mice included here are only females from the (Cuomo et al. 2023) study and the current 14-day study of lead acetate exposure in water. Animals exposed for 28-days (n = 3) are in green and animals exposed for 14-days (n = 5) are in orange. All samples are transformed log (y+1).

### Inter-strain variability in metal accumulation for Pb, Cr, and Cd

Mice exposed to Pb displayed the same rank order of Pb accumulation as observed in previous studies with CC006 accumulating the highest Pb in the blood, followed by CC035, CC057 and lastly CC019 (Fig. 3a). CC006 females accumulated higher amounts of Pb in the blood than every other pairwise comparison to females of other strains (6 vs 35, 6 vs 57, 6 vs 19).

**Fig. 3.**
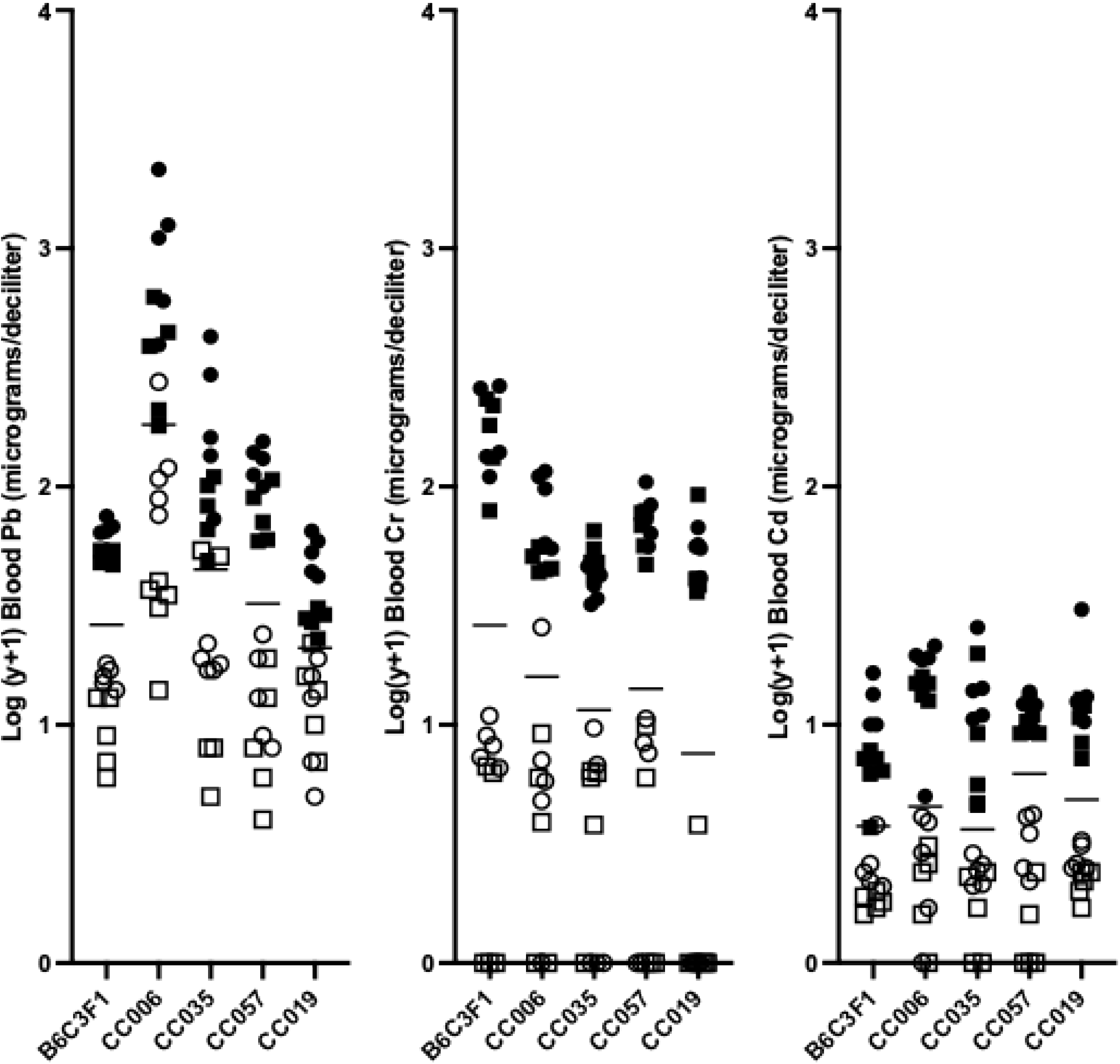
Log transformation (y+1) of blood metal levels measured in micrograms per deciliter. a) Pb; b) Cr; and c) Cd blood levels. Filled symbols are mice on high dose and open symbols are mice on low dose. Each strain group has males and females of the low and high dose represented. Circles are female, squares are male.

Mice exposed to Cr did not display the same strain rank accumulation patterns as mice exposed to Pb (Fig. 3b). Cr accumulation was significantly different between CC019 and B6C3F1 on low dose (p = 0.0206) after Tukey correction. B6C3F1 mice on high dose Cr consistently accumulated higher amounts of blood Cr compared to every CC strain (p = <0.0001 for each pairwise comparison).

Mice exposed to Cd also displayed different strain rankings than the Pb exposed mice (Fig. 3c). ANOVA and Tukey comparisons showed no significant differences between the strain means for low dose Cd. However, CC006 was significantly higher than B6C3F1 at high dose (p = 0.0244).

### Sex Differences in metal accumulation in blood

Sex differences in the strains were observed in the 14-day exposure. The largest difference was observed in the CC006 strain exposed to Pb. Females of this strain accumulated higher amounts of Pb in blood than males (p = 0.0078, low dose; p = 0.0041, high) dose (Fig. 4a). Additionally, Female CC035 mice exposed to high dose lead accumulated more than males (p = 0.0357).

**Fig. 4.**
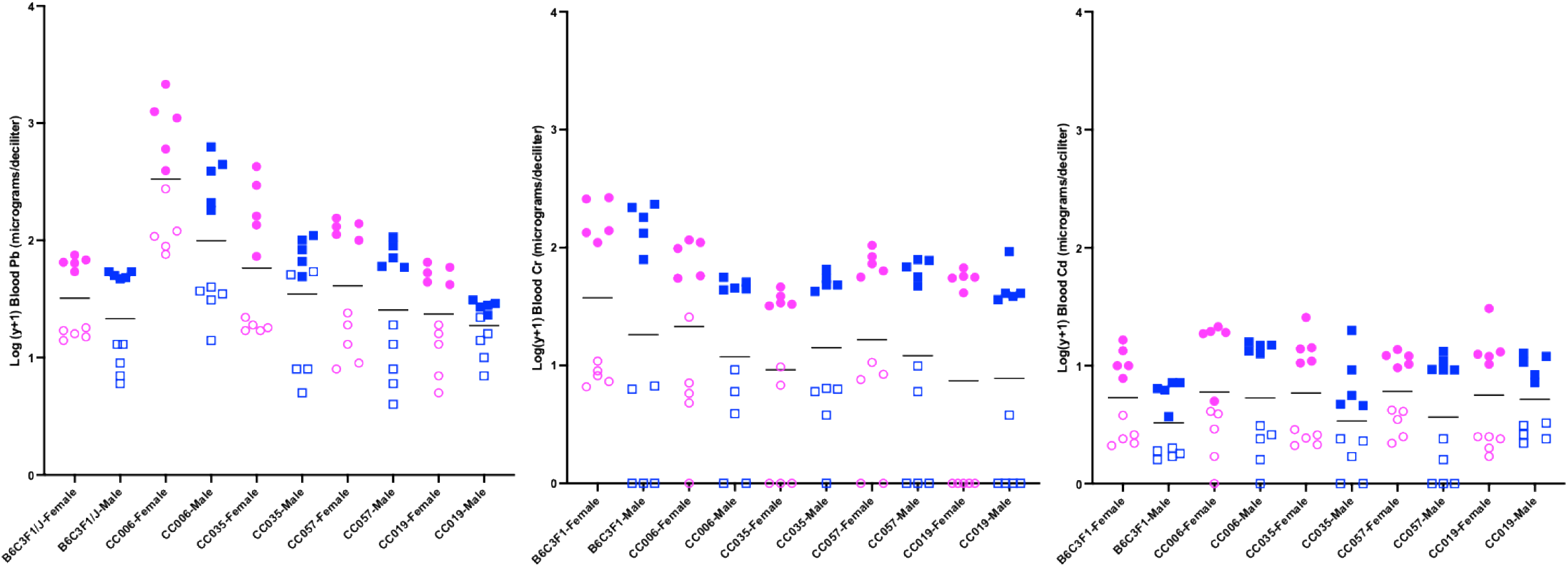
For all figures group metal levels are measured in micrograms per deciliter. a) Pb; b) Cr; and c) Cd blood levels. Blue boxes are males. Pink circles are females. Filled shapes are animals exposed to a high dose and unfilled shapes are animals exposed to a low dose.

Sex differences were not observed for Cr exposed mice at either dose (Fig. 4b). Similalry, minimal sex difference were observed in Cd exposed mice (Fig. 4c), with the only sex difference being between CC057 males and females on the low dose (p = 0.0024).

### Partitioning sources of variation

The effects of strain, sex, sex-by-strain interaction, water consumption, body weight and body weight change and their relative contribution to variance in blood Pb, Cr, or Cd was evaluated (Fig. 5). Variance partitioning showed that dose is the largest contributor to variance observed in all metal exposure paradigms. Addressing each metal exposure as a population with all animals grouped together for analysis, strain has the second largest effect for Pb exposure, but not for Cr or Cd exposure. The small effect of strain found in Cr and Cd exposed cohorts suggests that there may be other CC strains that would better represent the population phenotypic variability for Cr and Cd. This outcome suggests that genetics has a larger effect on Pb blood accumulation than it does for Cr or Cd in this subset of the CC population. A full summary of effect size for each metal, at each dose, and population wide are available in Supplemental Table 1.

**Fig. 5.**
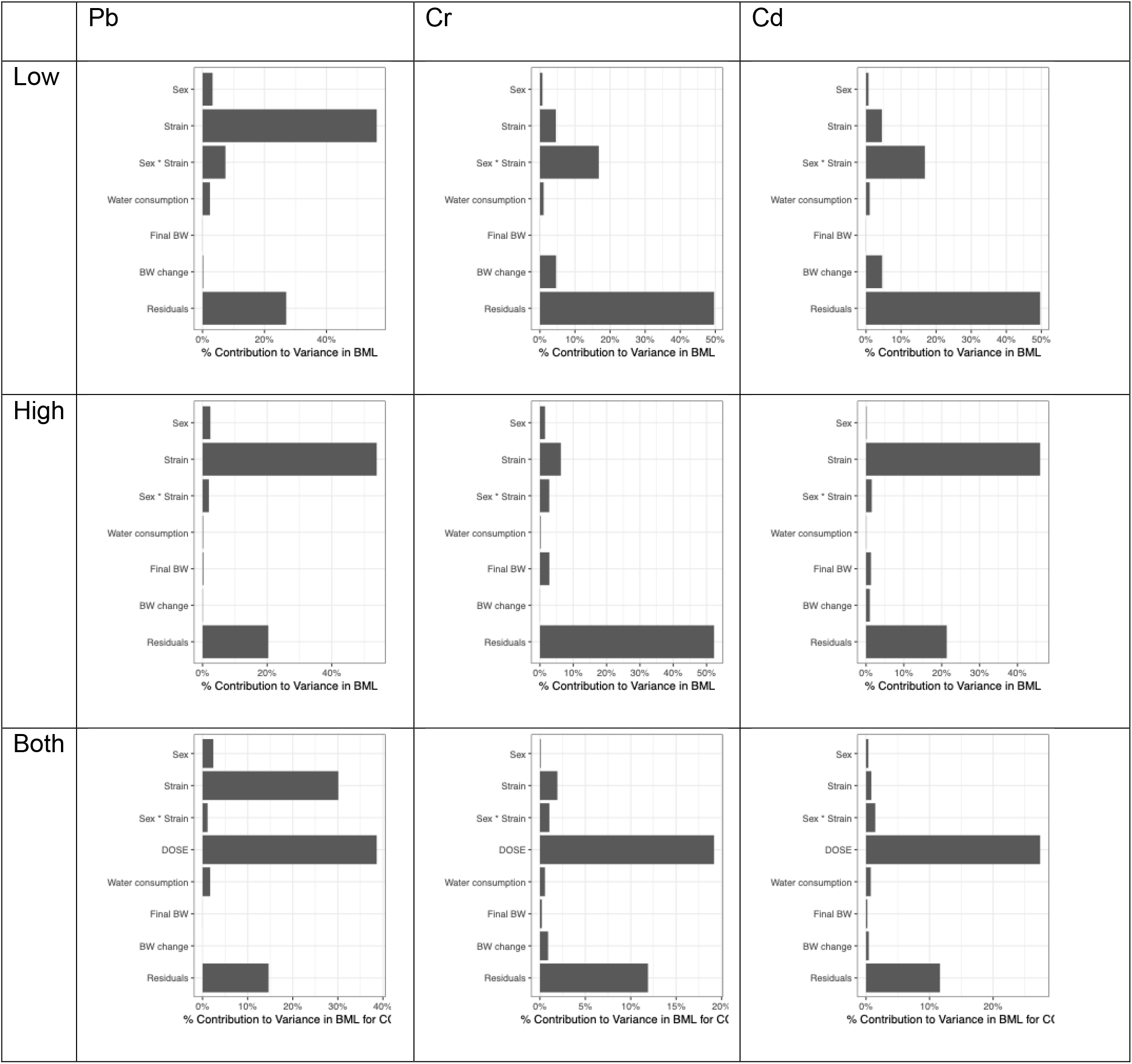
Partitioning of variance results for each metal using linear modeling in r to estimate eta squared.

## Discussion

Pb-blood accumulation rank order was the same between 28-day and 14-day exposure showing that shorter exposure paradigms yield similar results for blood accumulation as the dependent variable. A limitation to this conclusion is that while 28-day and 14-day rank accumulation is similar, these results are likely not extrapolatable to long term chronic exposure to Pb. However, strain rank accumulation for the three metals differed indicating that even a small panel of strains will not suffice for estimating variation across all toxicants.

Similar to the current study, sex difference observed in Pb exposed animals have been documented in the literature. In Pb inhalation studies conducted on outbred CD-1 mice it was found that female mice accumulated more Pb into the lungs than did male mice (González Rendón et al. 2018). Sex differences have also been recorded in blood Pb level accumulation in humanized Apolipoprotein E3 and E4 knock-in (ApoE3KI and ApoE4-KI) mice (Engstrom et al. 2017). Sex differences were not observed as significant contributors to variance in blood metal levels for Cr or Cd exposures.

The strains used in this study were chosen based on their response in blood level to Pb. Use of these strains for other metals does not capture the same range of blood metal level response. For Cd and Cr specifically, this suggests the full phenotypic range of blood accumulation may not be captured with a limited number of strains. Each metal has a different response profile when attributing causal factors of variance in blood metal levels. For example, sex is a small contributing factor to blood Pb level but not for Cr or Cd blood levels.

To adequately capture the population variability for each metal and likely other toxicants, a larger panel of strains will be required. In addition, residual variance might be reduced by including biological factors such as metal excretion or indicators of liver and kidney function.

The divergence in strain metal accumulation ranking supports the necessity of multiple strains in toxicological profiling which is currently not done by using of a single F1 hybrid strain in formal toxicity testing paradigms. Since each metal has a different strain response profile, the incorporation of multiple strains elucidates different susceptibilities to the three metal toxicants investigated. This finding supports the need for a large panel strains when formulating policy and regulation to estimate population response, rather than a single strain that may over or under protect certain members of a population.

## Conclusions

The well-known adage “The dose makes the poison” for the basis of toxicology and pharmacology is more accurately stated “The dose primarily makes the poison” since factors such as sex, strain, and behaviors like water consumption can also contribute. The sex difference observed in Pb exposed animals show that in further studies using both male and female mice should be examined when determining genetic susceptibility to heavy metals. It is also important to consider the use of larger panels of strains to capture population variability in susceptibility to these metals and likely other toxicants. These findings help inform development of more reliable toxicity ranges by experimentally modeling the genetic diversity of the human population.

## Acknowledgements

We thank the Texas A&M Institute for Genome Sciences and Society for access to equipment for metabolic chambers and MRI portions of experiments.

